# Phonological decoding ability is associated with fiber density of the left arcuate fasciculus longitudinally across reading development

**DOI:** 10.1101/2024.09.24.614799

**Authors:** Meaghan V. Perdue, Bryce L. Geeraert, Kathryn Y. Manning, Deborah Dewey, Catherine Lebel

**Author notes:** Corresponding Author: Meaghan Perdue, 28 Oki Drive NW, Calgary, Alberta, Canada, T3B 6A8.

## Abstract

Numerous studies have linked reading ability to white matter microstructure using diffusion tensor imaging, but recent large studies have failed to show consistency. Fiber-specific diffusion-weighted magnetic resonance imaging (dMRI) models offer enhanced precision to measure specific features of white matter structure and may be more sensitive to individual differences in reading skills. However, fiber-specific models have not yet been applied to examine associations between reading ability and white matter microstructure over the course of reading acquisition. In this accelerated longitudinal study, we applied constrained spherical deconvolution (CSD) and fiber-specific modelling to characterize developmental changes in fiber density of key white matter tracts of the reading network bilaterally, and investigated associations between tract-wise fiber density and children’s phonological decoding abilities. Fiber density was measured from ages 2-13 years, and decoding ability (pseudoword reading) was assessed at ages 6 years and older. Higher decoding ability was associated with greater fiber density in the left arcuate fasciculus, and effects remained consistent over time. Follow-up analysis revealed that asymmetry changes in the arcuate fasciculus were moderated by decoding ability: good decoders showed leftward asymmetry from early childhood onward, while poorer decoders shifted toward leftward asymmetry over time. These results suggest that densely organized fibers in the left arcuate fasciculus serve as a foundation for the development of reading skills from the pre-reading stage through fluent reading. Ongoing developmental changes in fiber density and microstructural asymmetry throughout childhood may reflect a reciprocal relationship in which reading experience continues to refine and strengthen these pathways.

## 1. Introduction

Reading is critical to academic success and daily functioning beginning in childhood, and persistent reading difficulties can have detrimental consequences for academic performance, as well as social and mental well-being (Haft et al., 2016). Reading ability varies widely among individuals; those who fall at the lowest tail of the continuum may be diagnosed with a reading disability (i.e., dyslexia). A distributed network of brain regions support reading, including structures involved in sensory processing (auditory & visual), language, motor planning/control (articulation), and executive functions. The neural connections among these brain regions play an important role in reading; they coordinate functions required to convert text in its visual/orthographic form to phonological representations, meaning, and possibly verbal output. “Phonological decoding” refers to “sounding out” words by converting text (orthographic) input to its associated speech sounds (phonology). This ability is fundamental for the development of fluent reading (Hudson et al., 2009), and is thought to depend primarily on the neural connections linking occipito-temporal regions that process orthographic input to temporo-parietal and inferior frontal regions involved in phonological processing and speech output.

Prior research has shown associations between reading ability and diffusion properties measured with diffusion tensor imaging (DTI) in white matter of the reading network, including the arcuate fasciculus (AF) superior longitudinal fasciculus (SLF), inferior longitudinal fasciculus (ILF), and inferior fronto-occipital fasciculus (IFOF) (D’Mello & Gabrieli, 2018; Lebel et al., 2013; Vandermosten, Boets, Poelmans, et al., 2012; Vandermosten, Boets, Wouters, et al., 2012). There is also evidence that the trajectories of white matter tract development differ between good and poor readers, indicating that white matter plasticity may be associated with the development of reading skills (Lebel et al., 2019; Wang et al., 2017; Yeatman et al., 2012). However, recent evaluations of the literature question the reproducibility of previous findings and point out methodological limitations of DTI as a potential source of this heterogeneity (Huber et al., 2019; Koirala et al., 2021; Meisler & Gabrieli, 2022b; Moreau et al., 2018; Ramus et al., 2018). First, the tensor usually does not accurately reflect the complex organization of brain white matter, in which crossing, curving, and kissing fibers are highly prevalent (Jeurissen et al., 2013). Further, the metrics derived from DTI are not specific to microstructural features of white matter, which leads to vague or ambiguous interpretation (Raffelt et al., 2012; Raffelt, Tournier, et al., 2017). More advanced microstructural models with longitudinal data are needed to better characterize links between reading ability and white matter in the developing brain.

One such method is constrained spherical deconvolution (CSD; Calamuneri et al., 2018; Jeurissen et al., 2014). CSD is applied to dMRI data acquired with higher diffusion weightings and more directions than standard DTI to model multiple fiber populations within each voxel (modelled as a fiber orientation distribution function [FOD]) and more accurately represents complex fiber organization. It also separates intra- and extra-axonal water compartments to model specific white matter microstructural features, such as fiber density (Calamante et al., 2015; Raffelt et al., 2012; Raffelt, Tournier, et al., 2017). Apparent fiber density (henceforth, ‘fiber density’), fiber cross-section, and composite ‘fiber density and cross section’ metrics can be computed for independent fiber populations within each voxel (termed ‘fixels’).

Despite these advantages, few studies to date have applied CSD or other advanced dMRI models to investigate white matter microstructure in relation to reading ability (Geeraert et al., 2020; Huber et al., 2019; Koirala et al., 2021; Meisler & Gabrieli, 2022a; Vanderauwera et al., 2015; Zhao et al., 2016). Vanderauwera and colleagues (2015) compared DTI and spherical deconvolution models in a study examining associations between phonological awareness and microstructure of left hemisphere temporo-parietal tracts in pre-reading children (ages 5-6 years). They found significant associations between FA and phonological awareness in both the arcuate fasciculus and left projection tracts, but the spherical diffusion model showed enhanced specificity with significant associations between fiber-specific anisotropy and phonological awareness limited to the arcuate fasciculus. Zhao and colleagues (2016) applied spherical deconvolution to examine tract anisotropy and asymmetry in children with and without dyslexia. They reported higher fiber-specific anisotropy in the right SLF in children with dyslexia relative to controls; in addition, the children with dyslexia had reduced leftward microstructural asymmetry of the IFOF and elevated rightward microstructural asymmetry of the middle SLF branch that links the inferior parietal lobule to the precentral gyrus (SLF-II). These findings are consistent with alterations in the left hemisphere reading network and putative compensatory mechanisms in the right hemisphere. In another study, Huber and colleagues (2019) applied two advanced dMRI models to examine microstructure in the posterior corpus callosum in relation to reading ability, and reported reading-related associations with metrics related to tissue density. Geeraert and colleagues (2020) identified multivariate white matter microstructural components based on principal components analysis of dMRI (DTI and Neurite Orientation Density and Dispersion Imaging [NODDI; Zhang et al., 2012] models) and myelin-specific imaging (magnetization transfer imaging and multicomponent relaxometry). However, the white matter components were not significantly associated with reading ability in a sample of typically developing children and adolescents. Together, these studies illustrate the potential utility of advanced dMRI models in studies of reading ability, but further research using larger samples and longitudinal designs is needed to fully characterize associations between reading and white matter microstructure.

Advanced dMRI models have been applied in two recent studies of reading in large samples drawn from the Healthy Brain Network Biobank (Alexander et al., 2017). Koirala and colleagues (2021) found that the neurite density index and orientation dispersion index were negatively correlated with both phonological processing and composite reading ability (timed word and pseudoword reading) in tracts throughout the brain; in contrast, DTI metrics showed limited relations to phonological processing and no associations with reading ability. Meisler and Gabrieli (2022a) applied the CSD model and fixel-based analysis to show widespread associations between composite reading ability (timed word and pseudoword reading) and fiber density and cross-section (a metric thought to reflect capacity of white matter tracts to efficiently transmit information), but found only modest associations between FA and phonological decoding ability in follow-up analyses examining timed word and pseudoword reading separately (Meisler & Gabrieli, 2022b). Together, these studies illustrate the enhanced sensitivity of non-tensor dMRI models to individual differences in reading ability. However, these findings were derived from a single dataset that was oversampled for children with/at risk for neurodevelopmental conditions, so it remains unclear whether these effects generalize to typically developing children. Furthermore, no study to date has examined associations between reading ability and fiber-specific microstructure longitudinally from pre-reading to fluent reading.

Prior studies from our lab have shown that leftward microstructural asymmetry of the AF (Reynolds et al., 2019) and multimodal gray and white matter components in the reading network (Manning et al., 2022), are associated with reading-related skills (phonological processing and speeded naming) in early childhood. However, associations between word-level reading abilities and fiber-specific microstructural properties have not yet been examined. In this study, we characterized the development of fiber density in reading-related white matter tracts over the course of reading development, and examined associations between fiber density and decoding ability in children learning to read English. We predicted that fiber density would increase with age, and that better decoding ability would be associated with greater fiber density and faster increases of fiber density in left hemisphere reading-related tracts, with strongest effects in dorsal tracts (AF and SLF) thought to support phonological decoding (Vandermosten, Boets, Poelmans, et al., 2012). We performed post-hoc analyses to test associations between microstructural asymmetry and reading ability in tracts that showed significant associations with decoding ability in the main analysis.

## 2. Methods

### 2.1 Participants

Participants were drawn from a large accelerated longitudinal study of brain development spanning early childhood-early adolescence (Reynolds et al., 2020). All participants were born full term (≥37 weeks gestation), and no participants had diagnosed neurological, genetic, or neurodevelopmental conditions upon enrollment. This study was approved by the conjoint health research ethics board (CHREB) at the University of Calgary (REB13-0020). Parents provided written informed consent and children provided verbal assent prior to participation.

Children who completed an assessment of reading ability at one or more study visits at ages 6 years and older and had good quality dMRI data (with b=2000 s/mm^2^) from at least one study visit were included in analysis (quality assurance procedures described below). Analyses included 280 dMRI scans from 66 participants (34 females, 32 males, scan age range = 2.41-12.92 years [mean = 7.23, SD=2.43], reading assessment age range = 6-12.71 years [mean = 8.5, SD=1.82]); see Figure 1.

**Figure 1.**
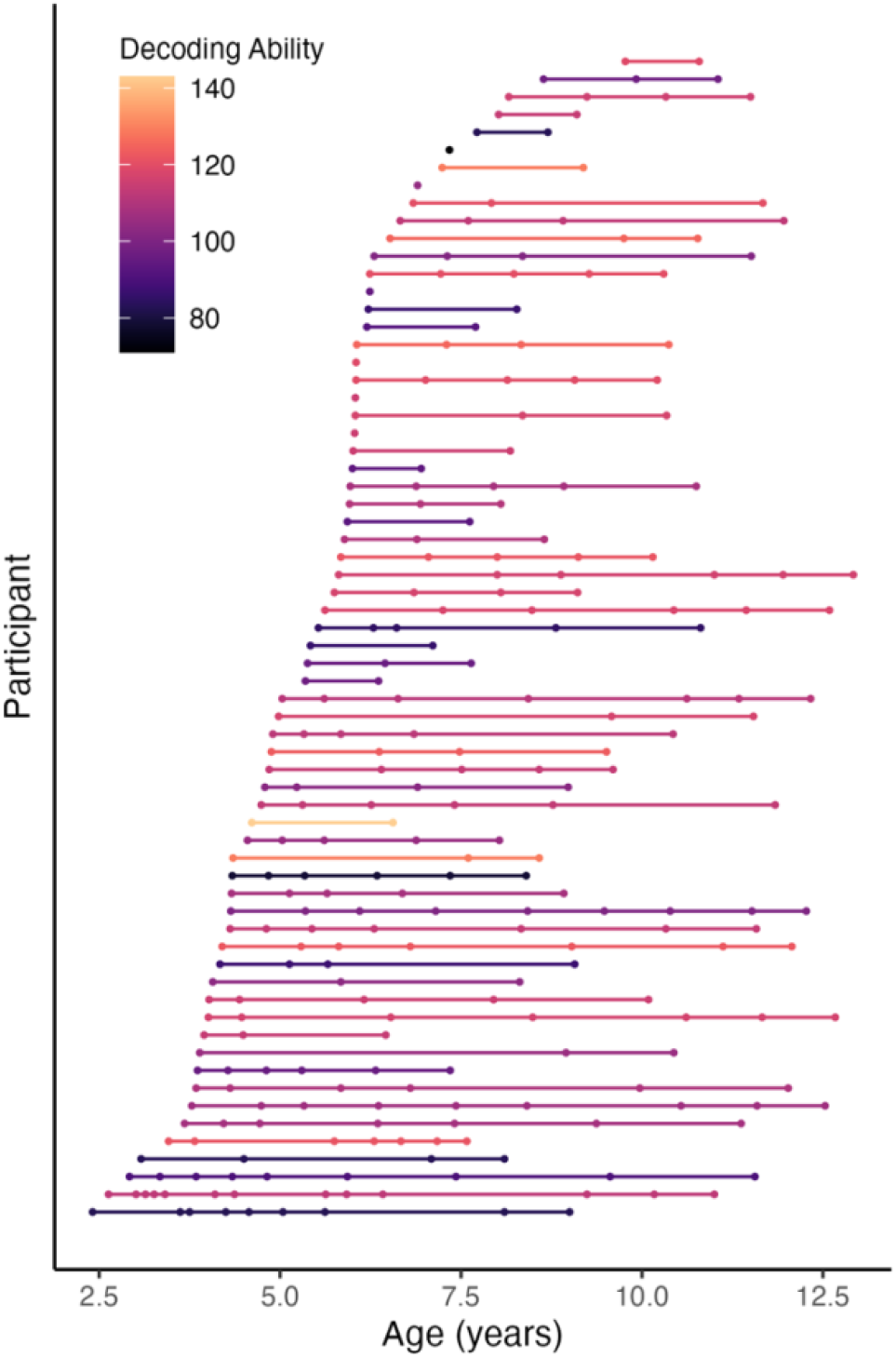
Age at scan acquisition by participant. Each scan is represented by a circle; each participant is shown in a different row with their scans connected by a straight line. Participants are colored by decoding ability (mean Word Attack standardized score across visits), with darker colors indicating poorer decoding ability.

### 2.2 dMRI acquisition and processing

dMRI data were acquired using a 3T GE MR750w MR system with a 32-channel head coil at the Alberta Children’s Hospital (Calgary, AB, Canada) without the use of sedation. dMRI data were acquired using a single shot spin echo EPI sequence (TR = 6750ms, TE = 97ms, 1.6 × 1.6 × 2.2 mm^3^ resolution [resampled on scanner to 0.78 × 0.78 × 2.2 mm], FOV = 20.0, 30 gradient encoding directions at b = 2000 s/mm^2^, and five interleaved volumes at b = 0 s/mm^2^. The imaging protocol also included acquisition of a separate dMRI sequence at b = 750 s/mm^2^, but these data were excluded from the present analyses due to the increased influence of extra-axonal water on apparent fiber density measures at low b-values (Calamante et al., 2015; Genc et al., 2020; Raffelt et al., 2012).

dMRI data were resampled to 2.2 × 2.2 × 2.2 mm^3^ isotropic voxels to match the slice thickness of the original acquisition, Gibb’s ringing correction was performed via MRTrix3 (v. 3.0.4; (Kellner et al., 2016; Tournier et al., 2019), and eddy current correction and motion correction with outlier replacement was performed using FSL’s *EDDY* (Andersson et al., 2016; Andersson & Sotiropoulos, 2016) via the *dwifslpreproc* function in MRTrix3. Preprocessed images were upsampled to 1 mm isotropic voxels, then binary brain masks were generated using *bet2* in FSL (Smith, 2002). dMRI data quality was assessed using the *EDDY QUAD* tool (Bastiani et al., 2019), and scans with average relative motion exceeding .7 mm and/or total outliers exceeding 10% were excluded from further processing (N=3). One additional scan was excluded because motion artifacts were apparent in the preprocessed image (these datasets were excluded prior to selection of data for analysis based on the criteria described above). Next, single-shell 3-tissue constrained spherical deconvolution (SS3T-CSD) was performed via MRtrix3Tissue (https://3Tissue.github.io, a fork of MRTrix3) using group average response functions. This method was chosen to optimize the fiber orientation distribution modelling by reducing the influence of gray matter-like and cerebrospinal fluid-like signal contributions and enhance sensitivity to intra-axonal signal by excluding a lower b-value shell (Dimond et al., 2020). Multi-tissue informed log-domain intensity normalization (*mtnormalise*; Dhollander et al., 2021; Raffelt, Dhollander, et al., 2017) was performed on the resulting fiber orientation distribution (FOD) maps for each tissue compartment. Fixel-based metrics were derived following the fixel-based analysis pipeline in MRtrix3 (Tournier et al., 2019); ‘fixel’ is defined as an individual fiber population within a voxel. A study-specific template was generated using FOD maps from 30 randomly selected sessions (all from different individuals). A template mask was generated as the intersection of all participants’ brain masks in template space and a corresponding fixel mask was generated. All participants’ FOD images were registered to template space, FOD lobes were segmented to estimate fixels, and apparent fiber density of each fixel was calculated. Each participants’ fixels were then reoriented in template space based on the local transformation at each voxel and assigned to template fixels.

### 2.3 Tractography

Automated tractography was performed on the FOD template via TractSeg (Wasserthal et al., 2018), and the following tracts were segmented bilaterally: arcuate fasciculi (AF), ventral superior longitudinal fasciculi (SLF-III; this branch was selected because of its terminations in temporo-parietal and inferior frontal cortices), inferior longitudinal fasciculi (ILF), and inferior fronto-occipital fasciculi (IFOF). Each tract was converted to a fixel map, which was thresholded to include only fixels with ≥5 streamlines passing through and then binarized to create a fixel mask. Mean fiber density of each tract was calculated as the mean fiber density of fixels included in the tract-specific fixel mask for each participant. Segmented tracts are shown in Figure 2. Code used for dMRI data processing and tractography is available here: https://github.com/developmental-neuroimaging-lab/mrtrix/tree/main/ss3tCSD_scripts

**Figure 2.**
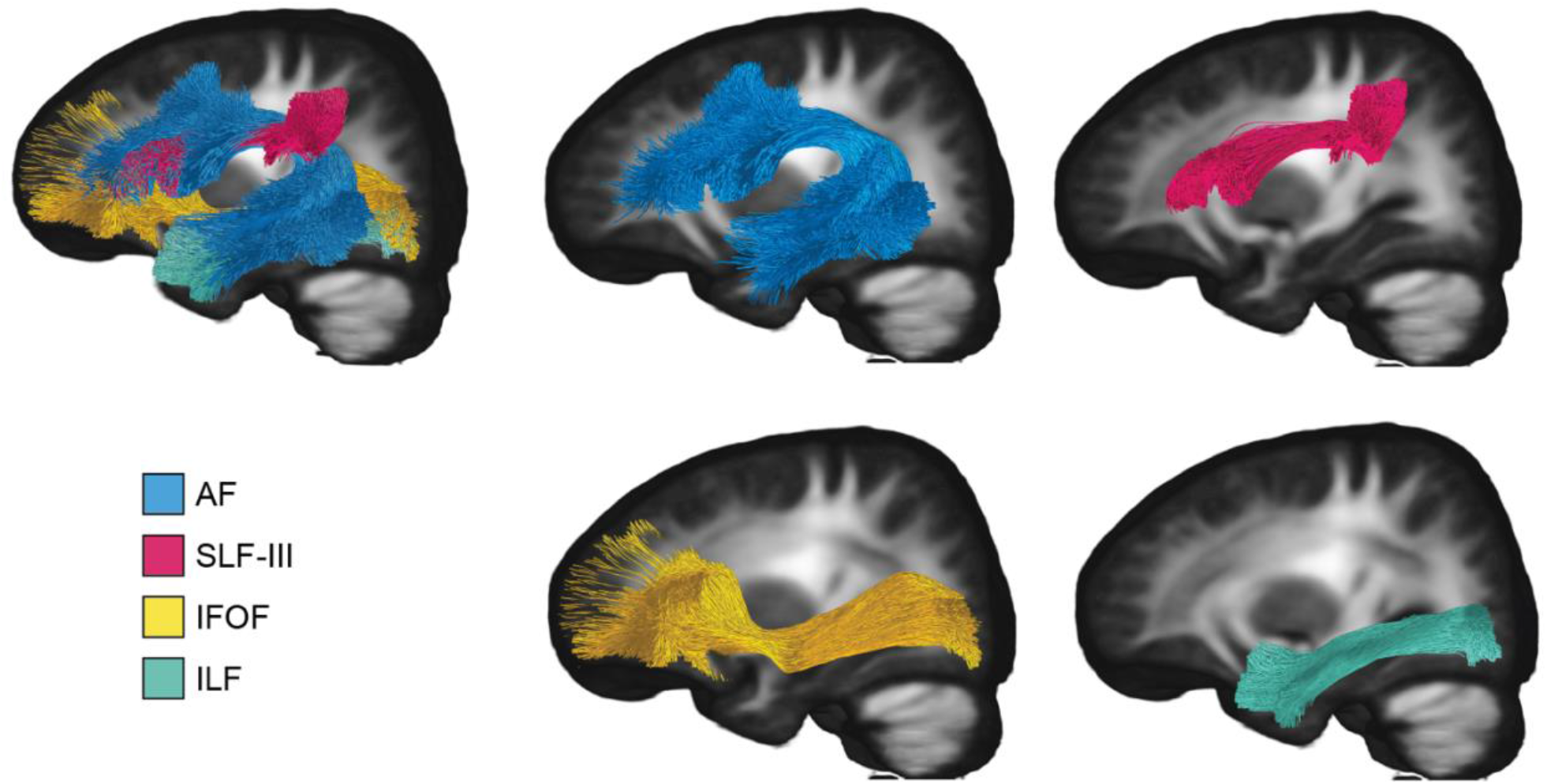
Reading-related tracts generated using TractSeg displayed on the study-specific template. Left hemisphere tracts shown for display, tracts were generated for both hemispheres. Blue=AF, Pink=SLF-III, Yellow=IFOF, Teal=ILF.

### 2.4 Reading Assessment

Reading ability was assessed at 6 years and older using the Woodcock Reading Mastery Tests 3^rd^ Edition (Woodcock, 2011) using the Word Identification and Word Attack subtests. We report descriptive statistics for both subtests in Supplementary Table 1. Our brain-behavior analyses focused on Word Attack because this assessment targets phonological decoding skills. It requires children to read aloud a series of pseudowords (spelled according to English conventions) of increasing difficulty. The test is not timed. We expected this subtest to be more sensitive to individual differences in phonological decoding abilities than Word Identification, which is an untimed test that requires the child to sight read English words. Raw scores were converted to age-normed standard scores, which have a population mean of 100 (SD = 15); higher scores indicate better performance. Standard scores of reading ability are expected to be stable over time in typically developing children, and are best described as a trait, rather than a variable skill (Woodcock, 2011; Yeatman et al., 2012). Accordingly, we calculated an average “decoding ability” score for each participant, which was the mean of their standard Word Attack scores across all available time points, excluding outlier scores (defined as a ≥30-point [2 SDs] difference between standardized scores of consecutive time points. Outlier scores were checked for scoring errors (none found) and removed for three individuals before computing their average decoding scores.

### 2.5 Statistical Analysis

We modelled the fiber density of each tract as a function of age and decoding ability using linear mixed effects regression with the *lme4* and *lmerTest* packages (Bates et al., 2015; Kuznetsova et al., 2017) in RStudio (RStudio Team, 2020). Mean fiber density values were scaled x100 to bring them to a closer scale to the other measures. We examined main effects of age and decoding ability on fiber density, and we tested whether rate of fiber density development was associated with decoding ability as the interaction between age and decoding score. Interaction terms were dropped from models when they did not show a significant effect. Models for each tract were defined as follows:

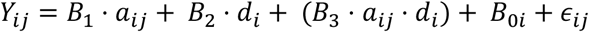

Where *Y*_*ij*_= fiber density of a given tract for the *i*th subject at the time of the *j*th scan, *B*_1_ = coefficient for age, *a*_*ij*_= *i*th subject’s age at time of *j*th scan, *B*_2_ = the coefficient for decoding ability, *d*_*i*_ = the *i*th subject’s decoding ability (time invariant), *B*_3_ = the coefficient for the interaction between age and decoding ability, *B*_0*i*_= subject-specific y-intercept, and ∈_*ij*_= random error.

False-discovery rate (FDR; Benjamini & Hochberg, 1995) was used to correct for multiple comparisons over eight tracts and two effects of interest (age and decoding ability). We report both uncorrected p-values and FDR-corrected q-values in the results.

Post-hoc analysis was performed to examine asymmetry of the AF as a function of age and decoding score. Asymmetry indices for fiber density of the AF were calculated as:

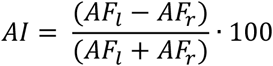

Where *AI* = asymmetry index of fiber density of the AF, *AF*_*l*_ = mean fiber density of the left AF, and *AF*_*r*_ = mean fiber density of the right AF. Greater values indicate stronger leftward asymmetry.

Asymmetry was modelled as a function of age and reading ability using the linear mixed effects model described above, substituting *AI* for *Y*.

## 3. Results

Descriptive statistics for age, reading ability, and fiber density are reported in Table 1.

**Table 1.** Descriptive statistics. Means reported across all included observations (*exception: Word Attack Mean Standardized Scores reported per individual participant).

|  | N obs. | Mean | SD | Min. | Max. |
| --- | --- | --- | --- | --- | --- |
| Age at MRI | 279 | 7.24 | 2.43 | 2.41 | 12.92 |
| Age at Reading Assessment | 159 | 8.50 | 1.82 | 6 | 12.71 |
| Word Attack (Decoding)<br>Mean Standardized Score | 66* | 107.66 | 14.21 | 71 | 143 |
| L. AF Fiber density | 279 | 0.45 | 0.02 | 0.39 | 0.51 |
| R. AF Fiber density | 279 | 0.44 | 0.02 | 0.38 | 0.50 |
| L. SLF Fiber density | 279 | 0.41 | 0.02 | 0.34 | 0.46 |
| R. SLF Fiber density | 279 | 0.41 | 0.03 | 0.32 | 0.46 |
| L. ILF Fiber density | 279 | 0.51 | 0.03 | 0.44 | 0.57 |
| R. ILF Fiber density | 279 | 0.49 | 0.03 | 0.41 | 0.55 |
| L. IFOF Fiber density | 279 | 0.55 | 0.03 | 0.49 | 0.60 |
| R. IFOF Fiber density | 279 | 0.53 | 0.02 | 0.46 | 0.59 |

### 3.1 Reading Ability

Overall, our sample had significantly higher Word Attack standard scores (mean=107.66, SD=14.21) relative to the normative sample (mean 100, SD = 15; *z*= 4.15, 95% CI of the mean [104.04, 111.28], *p*<.001); however, variability in reading scores was similar. Our sample included 6 children (9%) with poor decoding abilities (Word Attack Mean Standardized Scores *≤*85; at least 1 standard deviation below the mean of the norming sample), which is consistent with the population prevalence of reading disability (or developmental dyslexia) for which estimates range from ∼5-20% with recent estimates centering around 7% (Lyon et al., 2003; Peterson & Pennington, 2015; Yang et al., 2022). Descriptive statistics are reported in Table 1. Descriptive statistics for individual reading sub-tests are provided in Supplementary Materials Table S1.

### 3.2 Age-related changes in fiber density of reading network tracts

Fiber density increased significantly with age in all tracts, model statistics reported in Table 2.

**Table 2.**
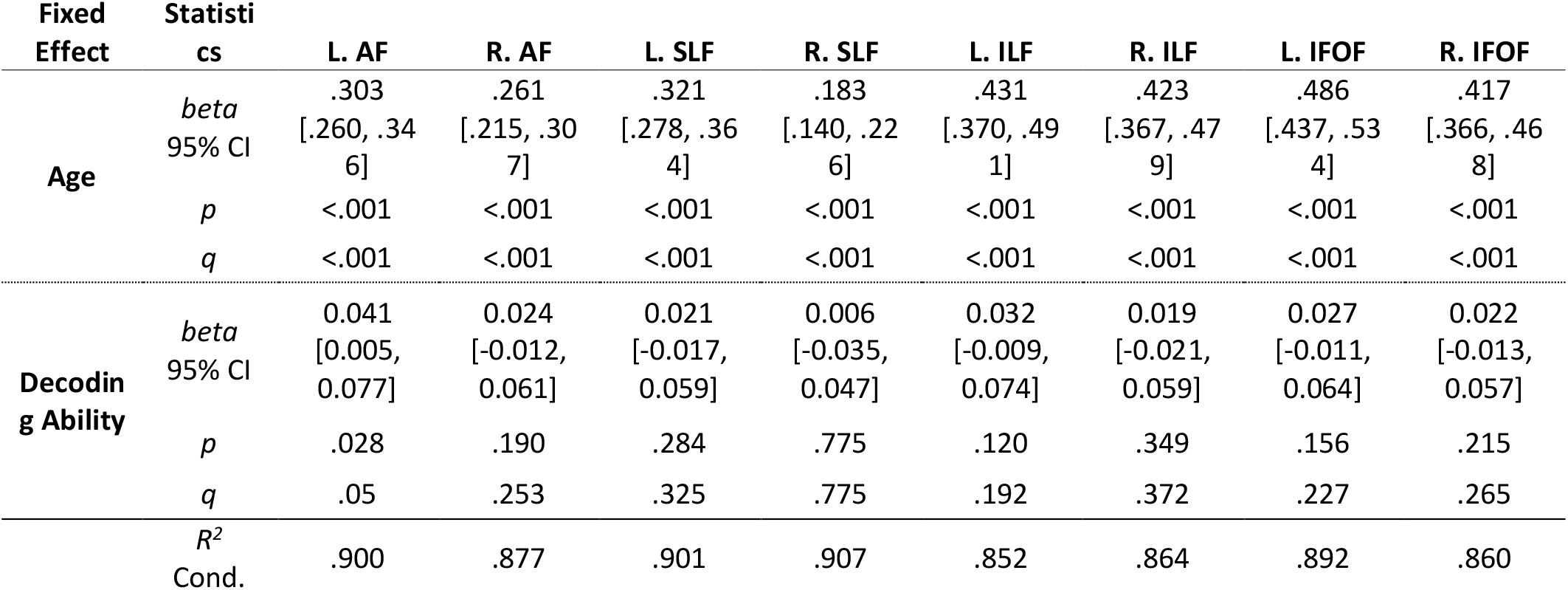
Model summaries showing age and decoding ability effects on fiber FD of each tract. Unstandardized beta values95% confidence intervals, uncorrected *p*-values, and FDR-corrected *q*-values are shown. R=Right, L=left.

### 3.3 Associations between fiber density and decoding ability

We observed a main effect of decoding ability on fiber density in the left AF, such that better decoders had higher initial fiber density that remained higher over time compared to poorer decoders (*beta*= .041, 95% CI[.005, .077], *p*=.028, *q*=.05) (Table 2; Figure 3). The interaction between age and decoding ability was not significant (*p*=.875), indicating that rate of fiber density increase did not vary based on decoding ability. Decoding ability was not significantly associated with fiber density in any other tract.

**Figure 3.**
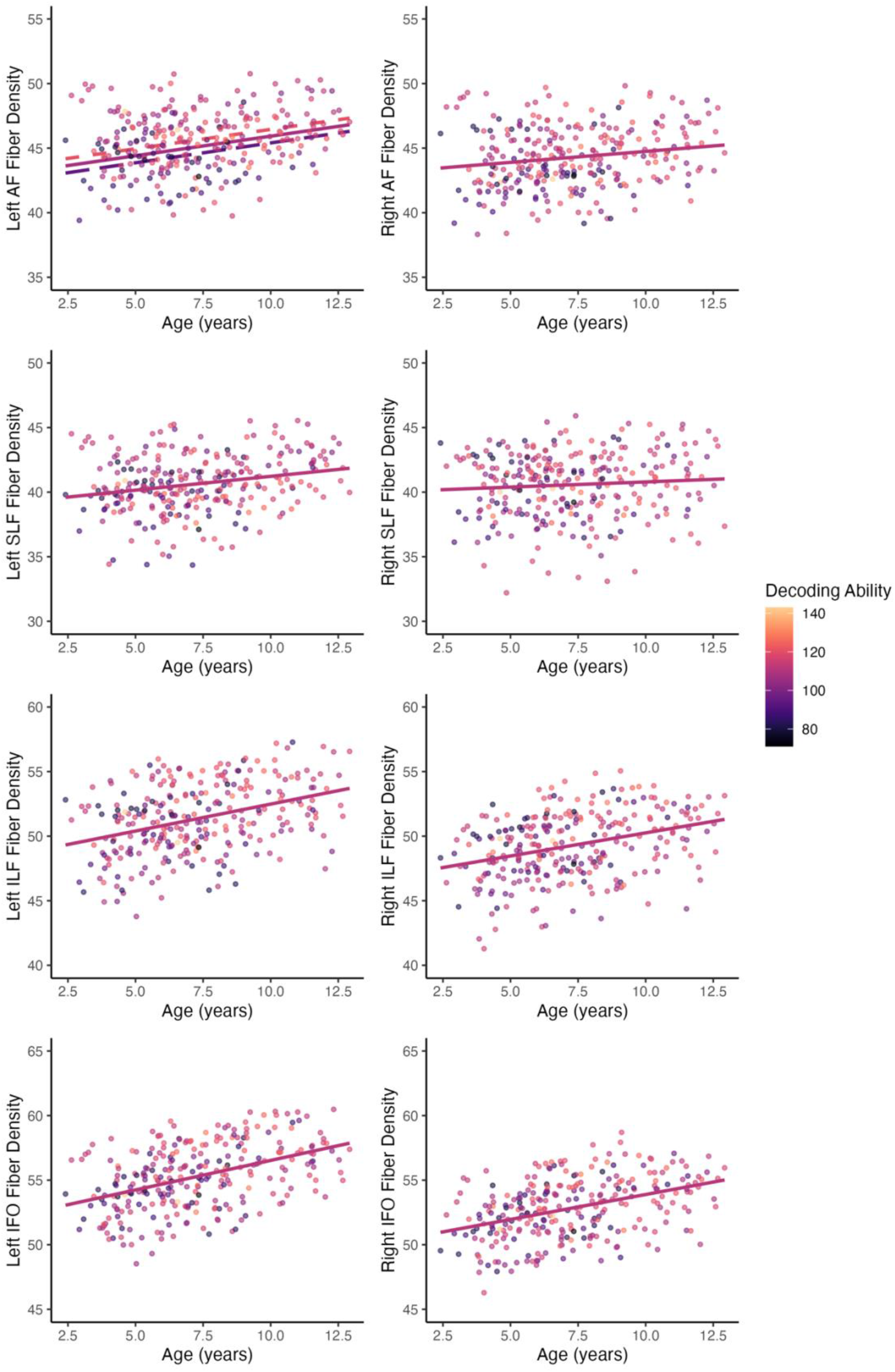
Scatterplots showing associations among age, fiber density and decoding ability. Dots are colored by decoding ability (mean Word Attack Standard Score per participant); darker colors indicate poorer decoding ability. Solid magenta lines show the overall fit for the age-fiber density association. Dashed fit lines for lower decoding ability (−1 SD from mean, dark purple long-dash) and higher decoding ability (+1 SD from mean, pink short-dash) are shown on the Left AF plot to illustrate main effect of decoding ability on fiber density.

### 3.4 Associations between fiber density asymmetry and decoding ability

We conducted post-hoc analysis to examine microstructural asymmetry of fiber density in the AF as a function of age and decoding ability. The model showed significant main effects of age and decoding ability on AF asymmetry, such that leftward asymmetry increased with age and decoding ability (*beta*_age_=.39, 95%CI[.03, .75], *p*_age_=.032; *beta*_decoding_=.04, 95%CI[0, .08], *p*_decoding_= .027). In addition, we observed a marginal interaction effect (*p*=.05) showing that better decoders had initial leftward asymmetry of the AF with little-to-no change over time, while poorer decoders had lower asymmetry of AF microstructure initially, which shifted toward more leftward asymmetry with age (Figure 4).

**Figure 4.**
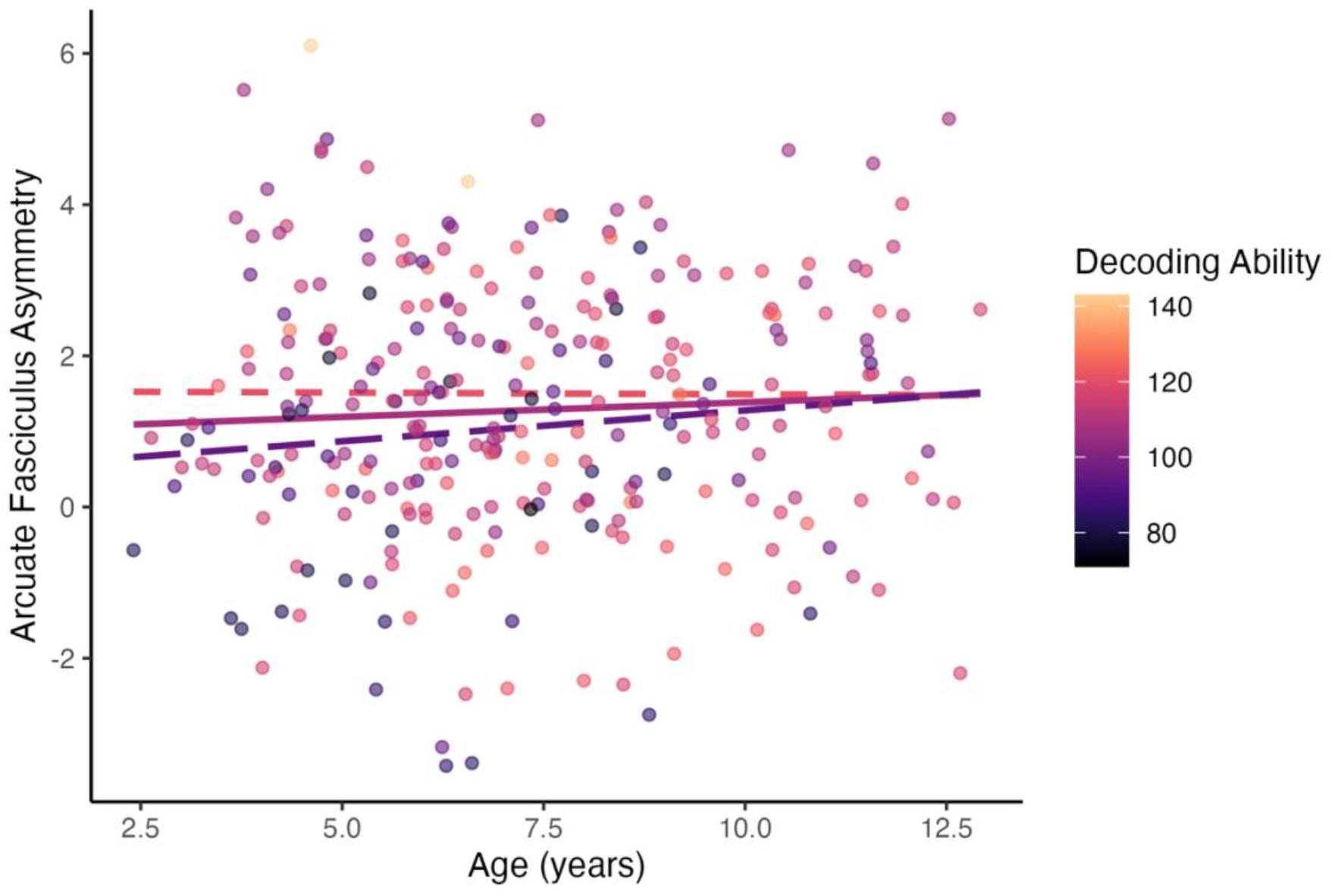
Scatterplot showing associations among age, asymmetry index of fiber density in the AF, and decoding ability. Higher asymmetry values indicate more leftward asymmetry. Dots are colored by decoding ability (mean Word Attack standard score per participant); darker colors indicate poorer decoding ability. Solid magenta line shows the mean fit for the age-asymmetry association. Dashed fit lines for lower decoding ability (−1 SD from mean, dark purple long-dash) and higher decoding ability (+1 SD from mean, pink short-dash) are shown to illustrate the moderating effect of decoding ability on asymmetry index, where children with lower decoding ability show increasing leftward asymmetry with age.

## 4. Discussion

In this study, we show a specific association between decoding ability and fiber density of the left AF across development of phonological decoding skills. The rate of fiber density development was not moderated by decoding ability, with good and poor decoders showing similar age-related changes. In contrast, microstructural asymmetry of the AF was associated with reading ability, and a marginal moderation effect by decoding ability was found (*p*=.05): good decoders had stable leftward asymmetry across the age range, while poorer decoders had increasing leftward asymmetry of AF fiber density with age. These findings distinguish fiber density as a key microstructural feature driving associations between white matter and reading abilities and highlight the left AF as an important tract for developing and sustaining reading skills.

The main effect linking left AF fiber density and phonological decoding ability (in the absence of an age-by-decoding ability interaction) indicates that better decoders had higher fiber density than poorer decoders across the whole age range. The effect we observed in the left AF corresponds to a recent study in which reading ability was strongly associated with a composite measure of fiber density and fiber bundle cross-sectional area in left temporo-parietal white matter (Meisler & Gabrieli, 2022a). Our finding is also consistent with prior evidence linking white matter microstructure in infancy and early childhood to later language and reading skills (Manning et al., 2022; Van Der Auwera et al., 2021; Zuk et al., 2021). Together with earlier findings, our results show that dense, coherent white matter connections in the left hemisphere reading network (here, the AF) in early childhood provide a neural foundation for learning to read and precede reading difficulties. The continued refinement of these fibers throughout childhood may reflect a reciprocal relationship with reading experience, as ongoing practice further strengthens white matter pathways.

Importantly, the present study enhances our understanding of the microstructural features that support skilled reading by showing a specific association between fiber density and decoding ability. This helps to clarify prior evidence that has shown associations between reading ability and tensor-based metrics such as FA (Catherine Lebel et al., 2013; Vandermosten, Boets, Poelmans, et al., 2012; Vandermosten, Boets, Wouters, et al., 2012; Wang et al., 2017; Yeatman et al., 2012). Our findings suggest that axon packing within the left AF is related to decoding skill; this architecture may reflect a greater capacity for efficient signalling between posterior and anterior reading network regions. The robustness of our CSD model to the influences of crossing fibers further enhances the specificity of these findings and provides confidence that fiber density, rather than the presence of crossing fibers, drives the observed brain-behavior associations.

Additional analyses showed that leftward asymmetry of AF fiber density increased with both age and decoding ability. A modest interaction effect indicated that children with better phonological decoding skills had leftward asymmetry of the AF in early childhood and showed little-to-no change with age, while poorer decoders exhibited increasing leftward asymmetry over time. Thus, along with overall greater fiber density of the left AF, early leftward asymmetry of the AF appears to predispose children to acquire strong reading skills. A previous study in an overlapping sample showed that macrostructural asymmetry of the AF is present by age two, but microstructural and functional asymmetry continue emerge into middle childhood (Reynolds et al., 2019). Here, we extend this finding to show that asymmetry of fiber density is established earlier in children who become strong readers. The leftward shift in AF asymmetry over childhood in poorer decoders may reflect experience-driven effects of learning to read on the micro-architecture of the left AF. Studies have shown mixed findings regarding changes in structural and functional asymmetry in response to reading intervention (Perdue et al., 2022), and further research is needed to disentangle practice-driven effects from maturational effects.

We did not observe significant associations with decoding ability in any other tracts examined, highlighting the regional specificity of these relationships to the left AF. The left AF connects key hubs of the reading network (i.e., occipito-temporal, temporo-parietal, and inferior frontal cortices), and decoding ability relies explicitly on orthographic-to-phonological mapping that is facilitated by the dorsal route *via* the temporo-parietal cortex (Jobard et al., 2003; Levy et al., 2009). Associations with ventral tracts may emerge when examining associations with reading skills that are more dependent on orthographic processing and rapid word recognition. For example, Vandermosten and colleagues (Vandermosten, Boets, Poelmans, et al., 2012) showed a dissociation between dorsal and ventral tracts in their associations with reading sub-skills in adults, such that dorsal tract microstructure was associated with phonological decoding while ventral tract microstructure was associated with orthographic processing.

The relatively small proportion of poor readers in our sample (n=6, 9%) may explain the modest effect sizes relative to case-control samples with matched numbers of children with good and poor reading abilities. The lack of poor readers in our study could also explain the absence of a moderation effect of reading ability on rates of fiber density development. Altered rates of tract development and/or effects in right hemisphere tracts could possibly emerge in studies targeting children with reading disabilities. For example, Huber and colleagues (2018) showed altered patterns of white matter microstructural development in the left AF, left ILF, and posterior corpus callosum of children with reading disabilities undergoing intensive reading intervention. Our study demonstrates the sensitivity and specificity of CSD tractography and fiber-specific microstructure measurement to individual differences in reading skills, supporting the extension of these methods to both typically developing readers and those with reading disabilities across all stages of reading acquisition.

## 5. Conclusion

Advanced dMRI modelling revealed a specific association between fiber density of the left AF and decoding ability over the course of reading acquisition. These findings highlight the left AF as a key tract supporting the acquisition of reading skills, especially phonological decoding. Fiber density of the left AF from early childhood (as young as age 2 years) through early adolescence was consistently associated with trait-level decoding abilities (measured at age 6 years and older), pointing to the architecture of this tract as a possible marker of later reading abilities. Ongoing tract microstructure changes and developmental shifts in asymmetry highlight the ongoing plasticity in reading network white matter. This protracted development points to the potential of the reading network to adapt in response to targeted reading interventions in children with reading difficulties.

## Data Statement

Tabular data and code for statistical analyses are available here:

https://osf.io/wmvgc/?view_only=f11afec2ea774e28b90495b4e9ccf0b4

Raw dMRI data is available upon request to the corresponding author.

Code for dMRI preprocessing, CSD, and tractography is available here: https://github.com/developmental-neuroimaging-lab/mrtrix

## Author Contributions (CReDIT Statement)

### Meaghan V. Perdue

Conceptualization, Methodology, Formal analysis, Writing – original draft, Writing –review & editing, Visualization. **Kathryn Y. Manning:** Methodology, Writing – review & editing. **Bryce L. Geeraert:** Methodology, Writing – review & editing. **Deborah Dewey:** Methodology, Writing – review & editing. **Catherine Lebel:** Conceptualization, Methodology, Writing – review & editing, Supervision, Funding acquisition.

## References

Alexander, L. M., Escalera, J., Ai, L., Andreotti, C., Febre, K., Mangone, A., Vega-Potler, N., Langer, N., Alexander, A., Kovacs, M., Litke, S., O’Hagan, B., Andersen, J., Bronstein, B., Bui, A., Bushey, M., Butler, H., Castagna, V., Camacho, N., … Milham, M. P. (2017). Data Descriptor: An open resource for transdiagnostic research in pediatric mental health and learning disorders. Scientific Data, 4, 1–26. 10.1038/sdata.2017.181

Andersson, J. L. R., Graham, M. S., Zsoldos, E., & Sotiropoulos, S. N. (2016). Incorporating outlier detection and replacement into a non-parametric framework for movement and distortion correction of diffusion MR images. NeuroImage, 141, 556–572. 10.1016/J.NEUROIMAGE.2016.06.058

Andersson, J. L. R., & Sotiropoulos, S. N. (2016). An integrated approach to correction for off-resonance effects and subject movement in diffusion MR imaging. NeuroImage, 125, 1063–1078. 10.1016/j.neuroimage.2015.10.019

Bastiani, M., Cottaar, M., Fitzgibbon, S. P., Suri, S., Alfaro-Almagro, F., Sotiropoulos, S. N., Jbabdi, S., & Andersson, J. L. R. (2019). Automated quality control for within and between studies diffusion MRI data using a non-parametric framework for movement and distortion correction. NeuroImage, 184(May 2018), 801–812. 10.1016/j.neuroimage.2018.09.073

Bates, D., Mächler, M., Bolker, B. M., & Walker, S. C. (2015). Fitting Linear Mixed-Effects Models Using lme4. Journal of Statistical Software, 067(i01). 10.18637/JSS.V067.I01

Benjamini, Y., & Hochberg, Y. (1995). Controlling the False Discovery Rate: A practical and powerful approach to multiple testing. Journal of the Royal Statistical Society, 57(1), 289–300.

Calamante, F., Smith, R. E., Tournier, J. D., Raffelt, D., & Connelly, A. (2015). Quantification of voxel-wise total fibre density: Investigating the problems associated with track-count mapping. NeuroImage, 117, 284–293. 10.1016/j.neuroimage.2015.05.070

Calamuneri, A., Arrigo, A., Mormina, E., Milardi, D., Cacciola, A., Chillemi, G., Marino, S., Gaeta, M., & Quartarone, A. (2018). White matter tissue quantification at low b-values within constrained spherical deconvolution framework. Frontiers in Neurology, 9(AUG), 1–14. 10.3389/fneur.2018.00716

D’Mello, A. M., & Gabrieli, J. D. E. (2018). Cognitive neuroscience of dyslexia. Language Speech and Hearing Services in Schools, 49(4), 798. 10.1044/2018_LSHSS-DYSLC-18-0020

Dhollander, T., Tabbara, R., Rosnarho-Tornstrand, J., Tournier, J.-D., Raffelt, D., & Connelly, A. (2021). Multi-tissue log-domain intensity and inhomogeneity normalisation for quantitative apparent fibre density. Proc. ISMRM, 29, 2472.

Dimond, D., Rohr, C. S., Smith, R. E., Dhollander, T., Cho, I., Lebel, C., Dewey, D., Connelly, A., & Bray, S. (2020). Early childhood development of white matter fiber density and morphology. NeuroImage, 210, 116552. 10.1016/j.neuroimage.2020.116552

Geeraert, B. L., Chamberland, M., Lebel, R. M., & Lebel, C. (2020). Multimodal principal component analysis to identify major features of white matter structure and links to reading. PLOS ONE, 15(8), e0233244. 10.1371/journal.pone.0233244

Genc, S., Tax, C. M. W., Parker, G. D., Jones, D. K., & Raven, E. P. (2020). Impact of b-value on estimates of apparent fibre density. Human Brain Mapping, 41, 2583–2595. 10.1002/hbm.24964

Haft, S. L., Myers, C. A., & Hoeft, F. (2016). Socio-emotional and cognitive resilience in children with reading disabilities. Current Opinion in Behavioral Sciences, 10, 133–141. 10.1016/j.cobeha.2016.06.005

Huber, E., Henriques, R. N., Owen, J. P., Rokem, A., & Yeatman, J. D. (2019). Applying microstructural models to understand the role of white matter in cognitive development. Developmental Cognitive Neuroscience, 36(June 2018), 100624. 10.1016/j.dcn.2019.100624

Hudson, R. F., Pullen, P. C., Lane, H. B., & Torgesen, J. K. (2009). The complex nature of reading fluency: A multidimensional view. Reading and Writing Quarterly, 25(1), 4–32. 10.1080/10573560802491208

Jeurissen, B., Leemans, A., Tournier, J. D., Jones, D. K., & Sijbers, J. (2013). Investigating the prevalence of complex fiber configurations in white matter tissue with diffusion magnetic resonance imaging. Human Brain Mapping, 34(11), 2747–2766. 10.1002/hbm.22099

Jeurissen, B., Tournier, J.-D. D., Dhollander, T., Connelly, A., & Sijbers, J. (2014). Multi-tissue constrained spherical deconvolution for improved analysis of multi-shell diffusion MRI data. NeuroImage, 103, 411– 426. 10.1016/j.neuroimage.2014.07.061

Jobard, G., Crivello, F., & Tzourio-Mazoyer, N. (2003). Evaluation of the dual route theory of reading: A metanalysis of 35 neuroimaging studies. NeuroImage, 20(2), 693–712. 10.1016/S1053-8119(03)00343-4

Kellner, E., Dhital, B., Kiselev, V. G., & Reisert, M. (2016). Gibbs-ringing artifact removal based on local subvoxel-shifts. Magnetic Resonance in Medicine, 76(5), 1574–1581. 10.1002/mrm.26054

Koirala, N., Perdue, M. V., Su, X., Grigorenko, E. L., & Landi, N. (2021). Neurite density and arborization is associated with reading skill and phonological processing in children. NeuroImage, 241, 118426. 10.1016/J.NEUROIMAGE.2021.118426

Kuznetsova, A., Brockhoff, P. B., & Christensen, R. H. B. (2017). lmerTest Package: Tests in Linear Mixed Effects Models. Journal of Statistical Software, 82(13), 1–26. 10.18637/JSS.V082.I13

Lebel, C., Benischek, A., Geeraert, B., Holahan, J., Shaywitz, S., Bakhshi, K., & Shaywitz, B. (2019). Developmental trajectories of white matter structure in children with and without reading impairments. Developmental Cognitive Neuroscience, 36. 10.1016/j.dcn.2019.100633

Lebel, C., Shaywitz, B., Holahan, J., Shaywitz, S., Marchione, K., & Beaulieu, C. (2013). Diffusion tensor imaging correlates of reading ability in dysfluent and non-impaired readers. Brain and Language, 125(2), 215–222. 10.1016/J.BANDL.2012.10.009

Levy, J., Pernet, C., Treserras, S., Boulanouar, K., Aubry, F., Démonet, J. F., & Celsis, P. (2009). Testing for the dual-route cascade reading model in the brain: An fMRI effective connectivity account of an efficient reading style. PLoS ONE, 4(8), 6675. 10.1371/journal.pone.0006675

Lyon, G. R., Shaywitz, S. E., & Shaywitz, B. A. (2003). Defining dyslexia, comorbidity, teachers’ knowledge of language and reading: A definition of dyslexia. Annals of Dyslexia, 53, 1–15. 10.1007/s11881-003-0001-9

Manning, K. Y., Reynolds, J. E., Long, X., Llera, A., Dewey, D., & Lebel, C. (2022). Multimodal brain features at 3 years of age and their relationship with pre-reading measures 1 year later. Frontiers in Human Neuroscience, 0, 594. 10.3389/FNHUM.2022.965602

Meisler, S. L., & Gabrieli, J. D. E. (2022a). Fiber-specific structural properties relate to reading skills in children and adolescents. ELife, 11, 1–28. 10.7554/ELIFE.82088

Meisler, S. L., & Gabrieli, J. D. E. (2022b). A large-scale investigation of white matter microstructural associations with reading ability. NeuroImage, 249, 118909. 10.1016/J.NEUROIMAGE.2022.118909

Moreau, D., Stonyer, J. E., McKay, N. S., & Waldie, K. E. (2018). No evidence for systematic white matter correlates of dyslexia: An Activation Likelihood Estimation meta-analysis. Brain Research, 1683, 36–47. 10.1016/j.brainres.2018.01.014

Peterson, R. L., & Pennington, B. F. (2015). Developmental Dyslexia. Annual Review of Clinical Psychology, 11(1), 283–307. 10.1146/annurev-clinpsy-032814-112842

Raffelt, D., Dhollander, T., Tournier, J.-D., Tabbara, R., Smith, R. E., Pierre, E., & Connelly, A. (2017). Bias field correction and intensity normalisation for quantitative analysis of apparent fibre density. Proc. Intl. Soc. Mag. Reson. Med, 25, 3541.

Raffelt, D., Tournier, J. D., Rose, S., Ridgway, G. R., Henderson, R., Crozier, S., Salvado, O., & Connelly, A. (2012). Apparent Fibre Density: A novel measure for the analysis of diffusion-weighted magnetic resonance images. NeuroImage, 59(4), 3976–3994. 10.1016/j.neuroimage.2011.10.045

Raffelt, D., Tournier, J. D., Smith, R. E., Vaughan, D. N., Jackson, G., Ridgway, G. R., & Connelly, A. (2017). Investigating white matter fibre density and morphology using fixel-based analysis. NeuroImage, 144, 58– 73. 10.1016/j.neuroimage.2016.09.029

Ramus, F., Altarelli, I., Jednoróg, K., Zhao, J., & Scotto di Covella, L. (2018). Neuroanatomy of developmental dyslexia: Pitfalls and promise. Neuroscience and Biobehavioral Reviews, 84, 434–452. 10.1016/j.neubiorev.2017.08.001

Reynolds, J. E., Long, X., Grohs, M. N., Dewey, D., & Lebel, C. (2019). Structural and functional asymmetry of the language network emerge in early childhood. Developmental Cognitive Neuroscience, 39(July), 100682. 10.1016/j.dcn.2019.100682

Reynolds, J. E., Long, X., Paniukov, D., Bagshawe, M., & Lebel, C. (2020). Calgary Preschool magnetic resonance imaging (MRI) dataset. Data in Brief, 29, 105224. 10.1016/J.DIB.2020.105224

RStudio Team. (2020). RStudio: Integrated Development Environment for R. http://www.rstudio.com/

Smith, S. M. (2002). Fast robust automated brain extraction. Human Brain Mapping, 17(3), 143–155. 10.1002/hbm.10062

Tournier, J. D., Smith, R., Raffelt, D., Tabbara, R., Dhollander, T., Pietsch, M., Christiaens, D., Jeurissen, B., Yeh, C. H., & Connelly, A. (2019). MRtrix3: A fast, flexible and open software framework for medical image processing and visualisation. In NeuroImage (Vol. 202, p. 116137). Academic Press Inc. 10.1016/j.neuroimage.2019.116137

Van Der Auwera, S., Vandermosten, M., Wouters, J., Ghesquière, P., & Vanderauwera, J. (2021). A three-time point longitudinal investigation of the arcuate fasciculus throughout reading acquisition in children developing dyslexia. NeuroImage, 237, 118087. 10.1016/j.neuroimage.2021.118087

Vanderauwera, J., Vandermosten, M., Dell’Acqua, F., Wouters, J., Ghesquière, P., Dell ‘acqua, F., Wouters, J., & Ghesquiè Re, P. (2015). Disentangling the relation between left temporoparietal white matter and reading: A spherical deconvolution tractography study. Human Brain Mapping, 36(8), 3273–3287. 10.1002/hbm.22848

Vandermosten, M., Boets, B., Poelmans, H., Sunaert, S., Wouters, J., Ghesquiè Re, P., Vandermosten, M., & Vesaliusstraat, A. (2012). A tractography study in dyslexia: neuroanatomic correlates of orthographic, phonological and speech processing. Brain, 135, 935–948. 10.1093/brain/awr363

Vandermosten, M., Boets, B., Wouters, J., & Ghesquière, P. (2012). A qualitative and quantitative review of diffusion tensor imaging studies in reading and dyslexia. Neuroscience & Biobehavioral Reviews, 36(6), 1532–1552. 10.1016/j.neubiorev.2012.04.002

Wang, Y., Mauer, M. V, Raney, T., Peysakhovich, B., Becker, B. L. C., Sliva, D. D., & Gaab, N. (2017). Development of tract-specific white matter pathways during early reading development in at-risk children and typical controls. Cerebral Cortex, 27, 2469–2485. 10.1093/cercor/bhw095

Wasserthal, J., Neher, P., & Maier-Hein, K. H. (2018). TractSeg - Fast and accurate white matter tract segmentation. NeuroImage, 183, 239–253. 10.1016/J.NEUROIMAGE.2018.07.070

Woodcock, R. W. (2011). Woodcock Reading Mastery Tests: WRMT-III. Pearson.

Yang, L. P., Li, C. B., Li, X. M., Zhai, M. M., Zhao, J., & Weng, X. C. (2022). Prevalence of developmental dyslexia in primary school children: a protocol for systematic review and meta-analysis. World Journal of Pediatrics, 18(12), 804–809. 10.1007/s12519-022-00572-y

Yeatman, J. D., Dougherty, R. F., Ben-Shachar, M., & Wandell, B. A. (2012). Development of white matter and reading skills. Proceedings of the National Academy of Sciences of the United States of America, 109(44). 10.1073/pnas.1206792109

Zhang, H., Schneider, T., Wheeler-Kingshott, C. A., & Alexander, D. C. (2012). NODDI: Practical in vivo neurite orientation dispersion and density imaging of the human brain. NeuroImage, 61(4), 1000–1016. 10.1016/j.neuroimage.2012.03.072

Zhao, J., Thiebaut de Schotten, M., Altarelli, I., Dubois, J., & Ramus, F. (2016). Altered hemispheric lateralization of white matter pathways in developmental dyslexia: Evidence from spherical deconvolution tractography. Cortex, 76, 51–62. 10.1016/j.cortex.2015.12.004

Zuk, J., Yu, X., Sanfilippo, J., Figuccio, M. J., Dunstan, J., Carruthers, C., Sideridis, G., Turesky, T. K., Gagoski, B., Grant, P. E., & Gaab, N. (2021). White matter in infancy is prospectively associated with language outcomes in kindergarten. Developmental Cognitive Neuroscience, 50. 10.1016/J.DCN.2021.100973/WHITE_MATTER_IN_INFANCY_IS_PROSPECTIVELY_ASSOCIATED_WITH_LANGUAGE_OUTCOMES_IN_KINDERGARTEN.PDF

