## Supplemental Table 1 for "Phonological decoding ability is associated with fiber density of the left arcuate fasciculus longitudinally across reading development"

**Supplementary Materials**

Table S1. Woodcock Reading Mastery Test Raw and Standardized Subtest Scores for Word Identification (Word ID) and Word Attack. Means reported for N obs. with the exception of Word Attack (Decoding) Mean Standardized Score, which is reported for N individuals.

|  | **N obs.** | **Mean** | **SD** | **Min.** | **Max.** |
| --- | --- | --- | --- | --- | --- |
| **Age at MRI** | 279 | 7.24 | 2.43 | 2.41 | 12.92 |
| **Age at Reading Assessment** | 159 | 8.50 | 1.82 | 6 | 12.71 |
| **Word ID Raw Score** | 217 | 20.28 | 14.56 | 0 | 46 |
| **Word Attack Raw Score** | 214 | 11.63 | 9.08 | 0 | 26 |
| **Word ID Standardized Score** | 166 | 112.30 | 17.58 | 63 | 145 |
| **Word Attack Standardized Score** | 161 | 108.89 | 14.13 | 71 | 143 |
| **Word Attack (Decoding)**  **Mean Standardized Score** | 66* | 107.66 | 14.21 | 71 | 143 |
